# A regional One Health approach to the risk of invasion by *Anopheles stephensi* in Mauritius

**DOI:** 10.1101/2023.12.05.570234

**Authors:** Diana P. Iyaloo, Sarah Zohdy, Ryan Carney, Varina Ramdonee Mosawa, Khouaildi B. Elahee, Nabiihah Munglee, Nilesh Latchooman, Surendra Puryag, Ambicadutt Bheecarry, Hemant Bhoobun, Harena Rasamoelina-Andriamanivo, Said Ahmed Bedja, Joseph Spear, Thierry Baldet, Tamar E. Carter

## Abstract

**Background:** *Anopheles stephensi* is an invasive malaria vector in Africa that threatens to put an additional 126 million people at risk of malaria if it continues to spread. The island nation of Mauritius is highly connected to Asia and Africa and is at risk of introduction due to this connectivity. For early detection of *An. stephensi,* the Vector Biology and Control Division under the Ministry of Health in Mauritius, leveraged a well-established *Aedes* program, as *An. stephensi* is known to share *Aedes* habitats. These efforts triggered multisectoral coordination and cascading benefits of integrated vector and One Health approaches.

**Methods:** Beginning June 2021, entomological surveys were conducted at points of entry (seaport, airport) and on ships transporting livestock in collaboration with the Civil Aviation Department, the Mauritian Port Authority and National Veterinary Services.

A total of 39, 18, 723 mosquito larval surveys were respectively conducted in the seaport, airport and other localities in Mauritius while 20, two and 26 adult mosquito surveys were respectively conducted in the seaport, airport and twenty-six animal points. Alongside adult mosquito surveys, surveillance of vectors of veterinary importance (e.g.- Culicoides spp.) was also carried out in collaboration with National Parks and Conservation Service and land owners.

**Results:** A total of 8,428 adult mosquitoes were collected and 1,844 larval habitats were positive for mosquitoes. All collected mosquitoes were morphologically identified and 151 Anopheles and 339 Aedes mosquitoes were also molecularly characterized. Mosquito species detected were Aedes albopictus, Anopheles arabiensis, An. coustani, An. merus, Culex quinquefasciatus, Cx. thalassius and Lutzia tigripes. Anopheles stephensi was not detected. The One Health approach was shared with the French Agricultural Research Centre for International Development (CIRAD), strengthening collaboration between Mauritius and Réunion Island on vector surveillance at entry points and insecticide resistance monitoring. The Indian Ocean Commission (IOC) was also alerted to the risk of An. stephensi, leading to regional efforts supporting trainings and development of a response strategy to An. stephensi bringing together stakeholders from Comoros, Madagascar, Mauritius, Réunion Island and Seychelles.

**Conclusions:** Mauritius is a model system showing how existing public health entomology capabilities can be used to enhance vector surveillance and control and create multisectoral networks to respond to any emerging public and veterinary health vector-borne disease threat.

**Author summary:** The malaria mosquito, *Anopheles stephensi*, is invasive in Africa where it threatens to put an additional 126 million people at risk of malaria if it continues to spread throughout the continent. The island nation of Mauritius is highly connected to Asia and Africa through maritime trade and therefore may be at risk of *An. stephensi* introduction and establishment. Mauritius implemented a One Health approach, enhancing entomological surveillance at entry points and collaborating across sectors (e.g. veterinary services, sea and air port authorities, national parks and conservation, communities, etc.) conducted extensive integrated vector surveillance, inspecting 85,071 larval habitats, and analyzing 8,428 adult mosquitoes morphologically and molecularly. Although *An. stephensi* was not detected, the initiative catalyzed and strengthened multisectoral partnerships nationally and across the Indian Ocean region member states (Comoros, Madagascar, Mauritius, Réunion Island and Seychelles). Leveraging the threat of *An. stephensi,* Mauritius exemplifies utilizing existing capabilities to create multisectoral networks for effective vector surveillance and response.

## 1. INTRODUCTION

*Anopheles stephensi* is an invasive malaria vector in Africa that threatens to expose an additional 126 million people to the risk of the disease and expand the malaria landscape from a predominantly rural to an equally urban disease (Sinka *et al*., 2020). This mosquito originates from Asia, and it is the main vector of malaria in the Indian Subcontinent (Sinka *et al*., 2011). Its first detection on the African continent in Djibouti in 2012, was made following a surprising outbreak of malaria at a time when Djibouti was approaching malaria elimination status (Faulde *et al.,* 2014). In the eight years following detection, malaria cases in Djibouti increased over 36-fold from <2,000 cases per year to over 75,000 confirmed cases and 300,000 suspected cases in 2020 (WMR, 2022). In 2016, *An. stephensi* was reported from eastern Ethiopia but it has since been detected across the country and, in 2022, was associated with a malaria outbreak in the urban centre of Dire Dawa (Tadesse *et al*., 2023). *Anopheles stephensi* has now been confirmed in eight countries in Africa: Djibouti (2012), Ethiopia (2016), Sudan (2016), Somalia (2019), Nigeria (2020), Kenya (2022), Eritrea (2022) and Ghana (2022). The potential impact of continued expansion has led to a WHO initiative to stop the spread of this invasive species (World Health Organization, 2022), and organizations like the US President’s Malaria Initiative have modified policy and guidance to ensure enhanced surveillance and rapid response to the invader (PMI, 2022).

The ecology of *Anopheles stephensi* appears different compared to African malaria vectors (*An. gambiae*, *An. coluzzii*, *An. arabiensis*, *An. funestus* to name the most important), not only in its proclivity for the use of artificial larval habitats like urban *Aedes* (Balkew *et al*., 2020), but its biting behavior appears to be unique as it is not captured by typical malaria vector surveillance methods like human landing catches. However, while information is available on the bio-ecology of *An. stephensi* in Asia, its area of origin, the limited knowledge about its bionomics in new African environments is a challenge considering the high ecological plasticity of the species, and therefore makes targeted detection and selection of vector control tools a challenge.

In addition to being reported as a unique *Anopheles*, *An. stephensi* has widely been reported to have associations with livestock, and bloodmeal analyses have indicated feeding preferences for livestock where they are present. For example, initial detections in Djibouti were first found in livestock quarantine settings (Faulde *et al*. 2014). In its invasive range, *An. stephensi* was also found to be so strongly associated with livestock that vector control efforts in Pakistan leveraged this association and utilized cattle permethrin dips as an effective tool to control the species (Rowland *et al*., 2001). In its invasive range in Africa, and Ethiopia in particular, studies thus far have also shown that one of the best *An. stephensi* adult collection methods is by mouth or backpack aspiration of resting mosquitoes from livestock structures (Balkew *et al*. 2021; Tadesse *et al*. 2023). Similarly, in Ethiopia the majority of identified bloodmeals are from livestock (Balkew *et al*. 2021; Carter *et al*., 2020); however, this could be due in part to the sampling bias of the livestock-focused adult collection method.

The adaptation of *An. stephensi* to urban breeding sites (small artificial water containers), its greater tolerance to cold than that of African malaria vectors, and its ability to persist year round, including throughout dry seasons, represents a major risk of modification to the malaria epidemiological landscape in Africa, specifically urbanization and a risk of upward extension to the highlands (Ethiopia, Kenya, Madagascar) where populations are not pre-immune to malaria leading to greater likelihood of epidemics (Samarasekera, 2022)

It is unknown how exactly *An. stephensi,* endemic to the Arabian Peninsula and South Asia, arrived on the African continent. In countries where *An. stephensi* was first detected within or close to a seaport (Djibouti, Sudan, and Somalia), the species was hypothesized to have arrived via maritime trade. Although that has been challenged by the notion that *Anopheles* eggs cannot remain viable in the absence of water and would not survive the journey, work from 1927 by Chalam *et al*. indicated that the species may be viable in the absence of water for up to 12 days, and when replicated more recently, data indicate that unlike other *Anopheles* species, *An. stephensi* eggs can remain in the absence of water in humid environments for up to three weeks (Leite et al. *in prep*). Genetic and genomic data also indicates that the port city of Djibouti and the sites connected to it by proximity, key railway and auto traffic routes (Carter *et al*. 2021, Samake *et al*. 2023), have established populations of *An. stephensi*, which supports the role of this port city in *An. stephensi* spread.

Of particular concern to Mauritius and several African countries are that shipping routes between Asia and Africa have amplified and shortened since the 2000s . This can increase the likelihood of gravid adult female mosquitoes resting in shelters such as containers to survive and lay eggs on arrival in port areas. The travel time of ships between Asia and Africa are compatible with the life expectancy of an adult mosquito (three weeks in this case).

A study examining maritime trade connectivity between *An. stephensi-*endemic countries and all seaports in Africa showed that maritime trade routes alone from 2011, 2016, and 2020 highlight the highest connectivity with Djibouti and Sudan, indicating that those may be the most likely locations for *An. stephensi* to have first invaded the continent (Ahn *et al*., 2023). The authors also describe an interactive network of connectivity to predict countries at risk of *An. stephensi* introduction. When coupled with habitat suitability predicting likelihood of establishment, Djibouti and Sudan remain the countries at highest risk (Ahn *et al*. 2023). Key goods identified along these maritime trade routes include livestock exportation and importation (Ahn *et al*. 2023).

Maritime trade routes highlighted that the small island nation of Mauritius is the third most likely country to have an *An. stephensi* introduction, based on trade routes alone (Ahn *et al*., 2023), and acts as a critical bridge and port-of-call between the Asian and African continents. Mauritius is a malaria elimination country since 1998 with no local cases of malaria, but with imported cases ranging between 12 and 54 cases every year (MoH, 2022) and residual populations of the historic vector *An. arabiensis* (Boussès *et al*., 2018). As a country with tourism as their third pillar, malaria elimination efforts have been rigorous and significant effort is made to maintain its disease-free status with focused emphasis on the early detection and treatment, focalized vector control interventions around confirmed cases and rapid response against introduced vectors to ensure that they do not become established (Tatarsky *et al*., 2011).The possibility of an introduction of *An. stephensi* may threaten the gains made in maintaining malaria elimination on this island nation. The Vector Biology and Control Division, a department under the Ministry of Health in Mauritius, has a well-established mosquito surveillance and control program across the island, with particular strengths against *Aedes,* especially *Aedes albopictus*, the vector of dengue fever. In late 2021, other mosquito collection methods were integrated into the program – including the use of mosquito adult traps, the establishment of artificial breeding sites and the consideration of animal transport or assembly points and the involvement of several stakeholders were sought to enhance the surveillance of *Anopheles,* which concurrently improved the monitoring system for other mosquito vectors. Therefore, the threat of *An. stephensi* provided an opportunity to catalyze multisectoral efforts to respond to this and future vector-borne public health threats.

Here, we describe the implementation of an enhanced vector surveillance system for the early detection of *An. stephensi.* These activities were launched and leveraged into an integrated vector management program, in alignment with the Global Vector Control Response 2017-2030 strategy of the World Health Assembly (WHO, 2017). We also described the necessity of multisectoral collaborations and the opportunities it has created for enhanced surveillance of all arthropod vectors of human and animal diseases, community engagement in vector surveillance, and a catalyst for the coordinated development and strengthening of a medical and veterinary entomology workforce across the south-east Indian Ocean islands (Mauritius, Comoros, Reunion, Madagascar, Seychelles), leveraging the strengths of each country.

Finally, we describe how the small island nation of Mauritius initiated multi-sectoral coordination to respond to the threat of *An. stephensi*, and highlight the cascading regional One Health benefits of this action thus far, including enhanced vector surveillance tools and approaches to inform public health decision making. The activities implemented by Mauritius and subsequently the Indian Ocean Commission (IOC) with other partners (CIRAD, IPM), serve as a model framework for how to leverage existing entomological public health capacity dedicated essentially to one system (*Aedes* mosquito and arbovirus surveillance) to enhance multi-vector surveillance and control and create multisectoral networks to prevent and respond to any emerging public health threats in the region.

## 2. MATERIALS AND METHODS

### 2.1 Study site

Mauritius is a small island nation (1865 km^2^) of 1,265,000 inhabitants located in the South West Indian Ocean, 890 km east of Madagascar (Fig. 1A). The island has two ports of entry – a sea port situated in the capital city of Port Louis and an airport in the southeast of the country, close to the locality of Plaine Magnien (Fig. 1B). The climate is mild tropical, consisting of a warm humid summer (November to April) and a cool dry winter (June to September). Mean temperature and mean annual rainfall are respectively 24.7^◦^ C and 1,344 mm during summer and 20.4^◦^ C and 666 mm during winter (MMS, 2012). Located in the South West Indian Ocean, Mauritius is particularly vulnerable to the impacts of climate change (pronounced due to the El Nino effect) which include rising temperatures and sea levels, coastal erosion, altered precipitation patterns, and an increase in extreme weather events such as flash floods and explosive cyclones (WHO, 2021). Mauritius has extensive maritime links with countries of the region and is at a high risk of being colonized by the invasive malaria vector *An. stephensi* (Ahn *et al*. 2023).

**Fig. 1:**
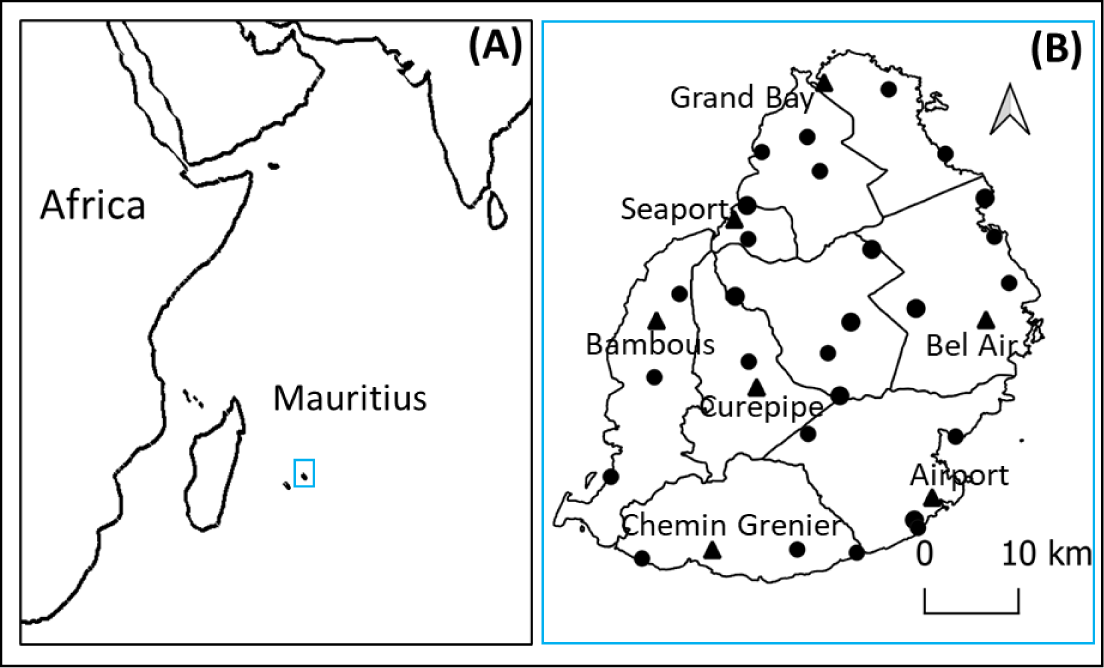
(**A)** Mauritius, an island in the Indian Ocean with (**B)** its two points of entry (seaport, airport) and five other sentinel sites (triangle) and 26 animal assembly points (dots) that were surveyed from June 2021 to September 2023.

### 2.2 Study design

#### 2.2.1 Larval mosquito survey

From June 2021 to March 2023, at least one larval survey was conducted every month in seven previously established sentinel sites (including the seaport and airport area) (Fig. 1B). Larval surveys were also conducted in randomly selected localities across the island. Since human population density is a major factor influencing mosquito vector populations and the risk of disease transmission, this parameter was used to estimate the relative frequency at which random surveys need to be carried out in each of the nine districts of the island. On a monthly basis, a pre-determined number of localities was therefore randomly selected from an exhaustive list of localities in each district.

During a larval survey, yards, plantations, vegetated areas and abandoned parcels within a delimited area, were systematically screened for the presence of stagnant water. All the larvae detected in small breeding sites were collected using pipettes and ladles. In larger water bodies (notably, drums, pails, buckets and basins), breeding sites were sampled using a ladle following the ‘Dipping method’ (WHO, 1975).

As such, respectively 39, 18, 113 and 610 mosquito surveys have been conducted in the seaport, airport, five other sentinel sites and randomly selected localities in Mauritius during which 5,230, 2,062, 14,343 and 78,571 potential outdoor breeding sites were inspected. Larvae were collected in 25 ml plastic tubes and brought back to the lab for species identification. *Anopheles* larvae were bred to the adult stage for species confirmation – approximately 50% of the larvae collected emerged into adults.

To investigate mosquito breeding habitat preferences at points of entry, commonly encountered breeding sites were classified into eight categories: (1) Basin (large concrete, fibre-glass or ground structures of >250 litres, used to store water for domestic purposes); (2) Drum, pail and bucket (large iron or plastic containers of 20-200 litres used to store water); (3) Discarded tyre; (4) Small container (mostly discarded waste containers of less than 50 cl, such as tins and plastic cups); (5) Flower pot (pots used for planting, including their saucers); (6) Bottle (discarded glass or plastic bottles); (7) Absorption pit, canal and drain; and (8) Puddles (puddles on roads, boats and plastic sheets).

#### 2.2.2 Adult mosquito survey

From October 2021 to September 2023, 20, two and 26 adult mosquito surveys were respectively conducted in the seaport, the airport and at twenty-six animal assembly points in Mauritius (Fig. 1B). During those surveys, adult mosquitoes were collected from 3 pm. to 8 am. the following day using BG Sentinel traps baited with BG Lure and 2 kg of dry ice. Mosquitoes collected were subsequently brought back to the laboratory for species identification.

#### 2.2.3 Larval sentinel surveillance at seaport

From July 2022 to April 2023, 38 artificial breeding sites - consisting of four 20-L concrete ground pools, 20 1-L plastic containers, eight car tyres and six 50-L plastic drums – were set up in the sea port area (Fig. 2). These sites were inspected on a weekly basis and larvae brought back to the laboratory for species identification. *Anopheles* larvae were reared to the adult stage for species confirmation.

**Fig. 2:**
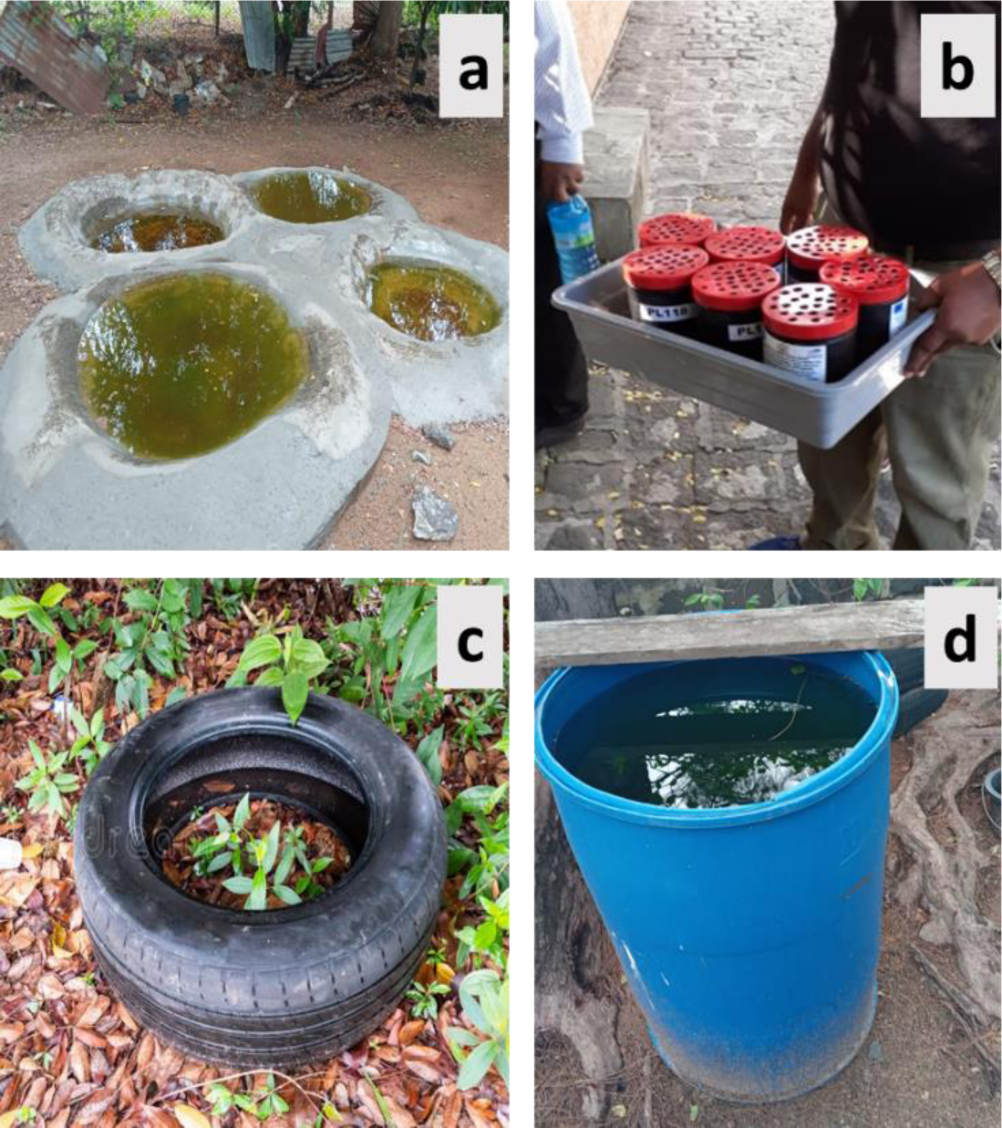
Examples of artificial breeding sites set up in the seaport area: **a** 20-L concrete ground pools; **b** 1-L plastic containers; **c** car tyre; **d** 50-L plastic drum

#### 2.2.3 Local stakeholders involved in mosquito surveys

Several areas where mosquito larval and adult surveys were carried out during this study are restricted or highly restricted zones, which required collaboration with several local stakeholders for ease of access. These stakeholders (summarized in Table 1), were involved in entomological surveillance at an early stage to ensure efficiency and sustainability.

**Table 1:** Mapping of local stakeholders involved in entomological surveillance in Mauritius.

| Location | Type and frequency of survey | Institution conducting surveys | Collaborators to access sites |
| --- | --- | --- | --- |
| Sea port area | Monthly larval survey<br>Weekly survey of artificial breeding sites<br>Snapshot adult mosquito survey | Vector Biology and Control Division | Mauritius Ports Authority |
|  |  |  | Port Health Office |
|  |  |  | Private companies |
|  |  |  | Community |
| Airport area | Monthly larval survey<br>Snapshot adult mosquito survey | Vector Biology and Control Division | Civil Aviation Department |
|  |  |  | Airport of Mauritius Ltd. |
|  |  |  | Airport Terminal Operations Ltd. |
|  |  |  | Plaisance Air Transport Services |
|  |  |  | Gound2Air Ltd |
|  |  |  | Plaisance<br>Meteorological<br>Station. |
|  |  |  | Beachcomber |
|  |  |  | Community |
| **Animal points** | ● Snapshot adult<br>mosquito survey | ● Vector Biology<br>and Control<br>Division<br>● Livestock and<br>Veterinary<br>Division | Livestock and<br>Veterinary Division |
|  |  |  | National Parks and<br>Conservation<br>Service |
|  |  |  | Private farmers |
| **Localities**<br>**island-wide** | ● Random larval survey | ● Vector Biology<br>and Control<br>Division | Institutions |
|  |  |  | Community |

### 2.3 Species identification

Excluding *Anopheles*, mosquito specimens collected during the surveys were identified morphologically under a stereomicroscope. Specimens were identified to the species level using the morphological keys of Edwards (1941), MacGregor (1927), Hamon (1953) and Coetzee *et al*. (2020). *Anopheles* larvae were reared to the adult stage for identification purposes using three methods: (1) morphological identification under a stereomicroscope at the VBCD laboratory using the morphological keys of MacGregor (1927) and Coetzee *et al*. (2020), (2) morphological analysis of smartphone photos of collected mosquitoes (larval and adult stages) using artificial intelligence system algorithms (Carney *et al*. in prep; Minakshi *et al*. 2020a,b; mosquitoID.org) at the University of South Florida, and (3) molecular analysis on a sub-sample of adult mosquitoes that were preserved on silica gel and sent to Baylor University.

DNA extractions of head and thorax were performed using the Qiagen DNeasy Blood and Tissue kit (Qiagen, Hilden, Germany). Extractions included samples morphologically identified as *An. arabiensis* (n=58) and *An. coustani* (n=10) across collection sites. To confirm species by molecular identification, portions of the internal transcribed spacer 2 (ITS2) locus and the mitochondrial cytochrome oxidase subunit I (COI) locus were PCR amplified and sequenced using protocols by Carter et al^1^. The ITS2 primer sequences used for *An. arabiensis* and *An. coustani* were 5.8SB (5ʹ-ATCACTCGGCTCGTGGATCG-3ʹ) and 28SB (5ʹ- ATGCTTAAATTTAGGGGGTAGTC-3ʹ). PCR amplifications were performed with the following temperature cycling: 95° C for 2 min, 30 cycles of 95° C at 30 s, 50° C at 30s, 72° C at 1min, and final extension of 72° at 5 min. The COI primer sequences used for all samples were LCO1490F (5ʹ-GGTCAACAAATCATAAAGATATTGG-3ʹ) and HCO2198R (5ʹ- TAAACTTCAGGGTGACCAAAAAATCA-3ʹ). PCR amplifications were performed with the following temperature cycling: 95° C for 1 min, 30 cycles of 95° C at 30 s, 48° C at 30 s, 72° C at 1min, and final extension of 72° at 10 min. For all samples, four microliters of PCR product were run on 2% agarose gel for 50 min at 100 V to confirm successful PCR. Amplified ITS2 and COI samples were sent out for commercial DNA Sanger sequencing.

DNA sequences were viewed and trimmed using CodonCode Aligner version 8 (CodonCode Corp., Centerville, MA, USA) with successfully sequenced ITS2 and COI sequences submitted as queries to the National Center for Biotechnology Information’s (NCBI) Basic Local Alignment Search Tool (BLAST) for final species confirmation. Sequences were used as the basis for further phylogenetic analysis. Sequences were submitted to NCBI Nucleotide database (Accession #). Alignments were generated using previously published sequences and those generated in this study with CodonCode and phylogenetic analysis was conducted using maximum-likelihood approach with RAxML [17] - Tamar. Final trees were annotated using Figtree [18] - Tamar.

The list of regional and international stakeholders that assisted in the identification of mosquito species during this study are summarized in Table 2.

**Table 2:** List of regional and international stakeholders involved in mosquito identification.

| Institution | Role |
| --- | --- |
| Centers for Disease Control and Prevention | ● Procurement of pinned specimens of adult mosquitoes to aid morphological identification<br>● Facilitate collaboration with other international collaborators |
| University of South Florida | ● Development and deployment of AI system to identify mosquito species through smartphone photos, targeting *An. stephensi* |
| Baylor University, Texas | ● Confirmation of mosquito species by PCR |
| NASA | ● Introduction to the Globe Mosquito Habitat Mapper, Citizen Science – a mobile application that help citizen survey and identify mosquito species in their surrounding |
| Indian Ocean Commission (IOC) | ● Organize training workshops in the morphological identification of *An. stephensi* for countries of the region (Mauritius, Reunion, Comoros, Madagascar, Seychelles) |
|  | • « The SEGA - One Health Network is the Indian Ocean Commission's arm in the field of public health, animal health and environmental health. It brings together more than 300 professionals from Member States' administrations and leading institutions. It is a platform for cooperation, promoting information sharing between Member States' health services and facilitating the mutualization of means and resources. Tangible results include consolidated surveillance, enhanced risk prevention, strengthened response capabilities, deployed technologies. » |
| French Agricultural Research Centre for International Development | • As a partner of the IOC, provide technical assistance in training countries of the region to the morphological identification of *An. stephensi* |
| Institut Pasteur Madagascar | • As a partner of the IOC, provide technical assistance in training countries of the region to the morphological identification of *An. stephensi* |

### 2.4 Habitat Data Analysis

To assess larval incidence, the container index (Focks, 2003) was calculated by dividing the number of habitats positive for mosquito larvae by the total number of habitats with water inspected during each survey. Breeding site preference for each mosquito species was evaluated by determining the frequency of their presence in the eight categories of potential breeding sites. Adult incidence was calculated as the total number of adults by species collected in one night.

## 3. RESULTS

### 3.1 Species identification

#### 3.1.1 Larval mosquito survey

In total, 780 larval surveys were conducted in 197 localities across the island from June 2021 to March 2023, during which 85,071 sites with water were inspected, and respectively 1,507, 258 and 78 sites were found positive for larvae of *Aedes albopictus*, *Culex quinquefasciatus* and *Anopheles gambiae sl* (presumed *An. arabiensis*) based on morphological identification. Sequencing of COI and ITS2 in 58 specimens confirmed *An. arabiensis* for the majority of the specimens and identified *An. merus* for two specimens (detailed below). Like in the other localities, *Ae. albopictus* was the dominant species in potential breeding sites at the seaport and airport, with 0.7 % and 0.8 % positive sites respectively. Sites occupancy by the two other mosquito species (*Cx. quinquefasciatus* and *An. arabiensis*) ranged from 0 to 0.2 % (Table 3). *Anopheles merus* was detected in only one site in Grand Bay in 2021. At both points of entry, small containers, flower pots, drums, pails and buckets were the most abundant sites containing water. However, although fewer in numbers, used tires were the most attractive breeding sites for *Ae. albopictus* at both the seaport and the airport (Tables 4 and 5). At the seaport, *Cx. quinquefasciatus* and *An. arabiensis* preferred to breed in basins (Table 4). Larvae of *Cx. quinquefasciatus* were not detected at the airport, while only one puddle (road pool) was found positive for *An. arabiensis* larvae (Table 5). No *An. stephensi* were detected during these larval mosquito surveys.

**Table 3:** Results of larval surveys conducted in seven sentinel localities at least once per month and in 190 randomly selected locations in Mauritius from June 2021 to March 2023.

| Locality | Type of survey | No. surveys | No. premises | No. sites | No. sites with water | No. sites with larvae by species (%) |  |  |
| --- | --- | --- | --- | --- | --- | --- | --- | --- |
|  |  |  |  |  |  | *Ae. albopictus* | *An. arabiensis* | *Cx. quinq* * |
| Seaport | Sentinel | 39 | 1559 | 5230 | 4042 | 30 (0.7) | 5 (0.1) | 8 (0.2) |
| Airport | Sentinel | 18 | 613 | 2062 | 1888 | 16 (0.8) | 1 (0.1) | 0 (0) |
| Bambous | Sentinel | 20 | 660 | 2367 | 2221 | 7 (0.3) | 3 (0.1) | 2 (0.1) |
| Bel Air | Sentinel | 23 | 1189 | 4013 | 3356 | 74 (2.2) | 4 (0.1) | 22 (0.7) |
| Chemin Grenier | Sentinel | 15 | 484 | 1756 | 1656 | 10 (0.6) | 2 (0.1) | 4 (0.2) |
| Curepipe | Sentinel | 27 | 641 | 2294 | 1933 | 9 (0.5) | 0 (0.0) | 2 (0.1) |
| Grand Bay** | Sentinel | 28 | 1332 | 3913 | 3301 | 41 (1.2) | 4 (0.1) | 4 (0.1) |
| 190 other localities | Random | 610 | 21243 | 78571 | 66674 | 1320 (2.0) | 58 (0.1) | 216 (0.3) |

\**Cx. quinq* = *Culex quinquefasciatus*; \*\* *Anopheles merus* was detected in only one site in Grand Bay

**Table 4:** Larval occupation of different categories of sites with water inspected during larval surveys carried out between June 2021 and March 2023 in the seaport area, Mauritius.

| Habitat type with water | No. surveyed | No. sites with larvae by species (%) |  |  |
| --- | --- | --- | --- | --- |
|  |  | *Ae. albopictus* | *An. arabiensis* | *Cx. quinq* * |
| Absorption pit, canal, drain | 358 | 0 (0.0) | 0 (0.0) | 0 (0.0) |
| Basin | 45 | 0 (0.0) | 4 (8.9) | 1 (2.2) |
| Bottles | 463 | 2 (0.4) | 1 (0.2) | 0 (0.0) |
| Discarded tyres | 86 | 8 (9.3) | 1 (1.6) | 0 (0.0) |
| Drum, pail, bucket | 655 | 6 (0.9) | 0 (0.0) | 0 (0.0) |
| Flower pot | 680 | 5 (0.7) | 0 (0.0) | 0 (0.0) |
| Puddle | 268 | 2 (0.7) | 0 (0.0) | 3 (1.1) |
| Small containers | 1487 | 7 (0.5) | 2 (0.1) | 1 (0.07) |

\* *Cx. quinq* = *Culex quinquefasciatus*

**Table 5:**
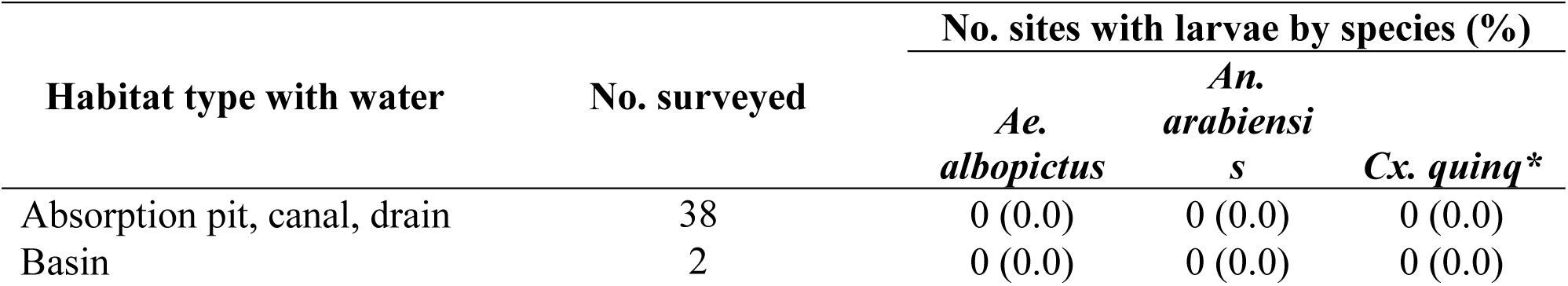
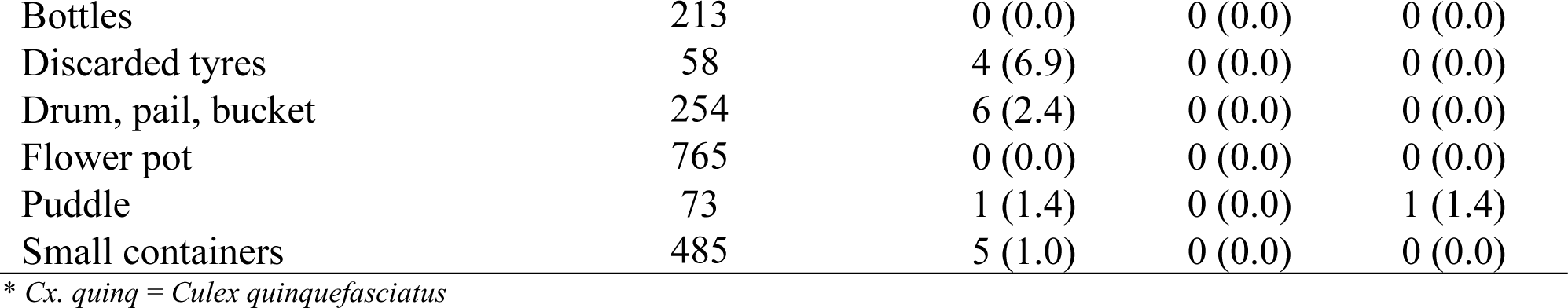
Larval occupation of sites with water inspected during larval surveys carried out between June 2021 and March 2023 in the airport area, Mauritius.

| Habitat type with water | No. surveyed | No. sites with larvae by species (%) |  |  |
| --- | --- | --- | --- | --- |
|  |  | *Ae. albopictus* | *An. arabiensis* | *Cx. quinq* * |
| Absorption pit, canal, drain | 38 | 0 (0.0) | 0 (0.0) | 0 (0.0) |
| Basin | 2 | 0 (0.0) | 0 (0.0) | 0 (0.0) |
| Bottles | 213 | 0 (0.0) | 0 (0.0) | 0 (0.0) |
| Discarded tyres | 58 | 4 (6.9) | 0 (0.0) | 0 (0.0) |
| Drum, pail, bucket | 254 | 6 (2.4) | 0 (0.0) | 0 (0.0) |
| Flower pot | 765 | 0 (0.0) | 0 (0.0) | 0 (0.0) |
| Puddle | 73 | 1 (1.4) | 0 (0.0) | 1 (1.4) |
| Small containers | 485 | 5 (1.0) | 0 (0.0) | 0 (0.0) |

\* *Cx. quinq* = *Culex quinquefasciatus*

#### 3.1.2 Adult mosquito survey

In total, 5,670 adult mosquitoes were collected by BG Sentinel traps at the seaport area during 20 nights of trapping as compared to 257 and 2,501 adult mosquitoes collected respectively at the airport and 26 animal assembly points during two and 26 nights of trapping from October 2021 to September 2023 (Table 6). Species collected were *Ae. albopictus*, *An. arabiensis*, *An. coustani, Cx. quinquefasciatus* and *Cx. thalassius.* The most abundant species was *Cx. quinquefasciatus* (3,754 in the seaport, 182 at the airport and 1,903 at animal assembly points) followed by *Ae. albopictus* (1,908 in the seaport, 75 at the airport and 525 at animal assembly points). No *An. stephensi* were detected during these adult mosquito surveys.

**Table 6:**
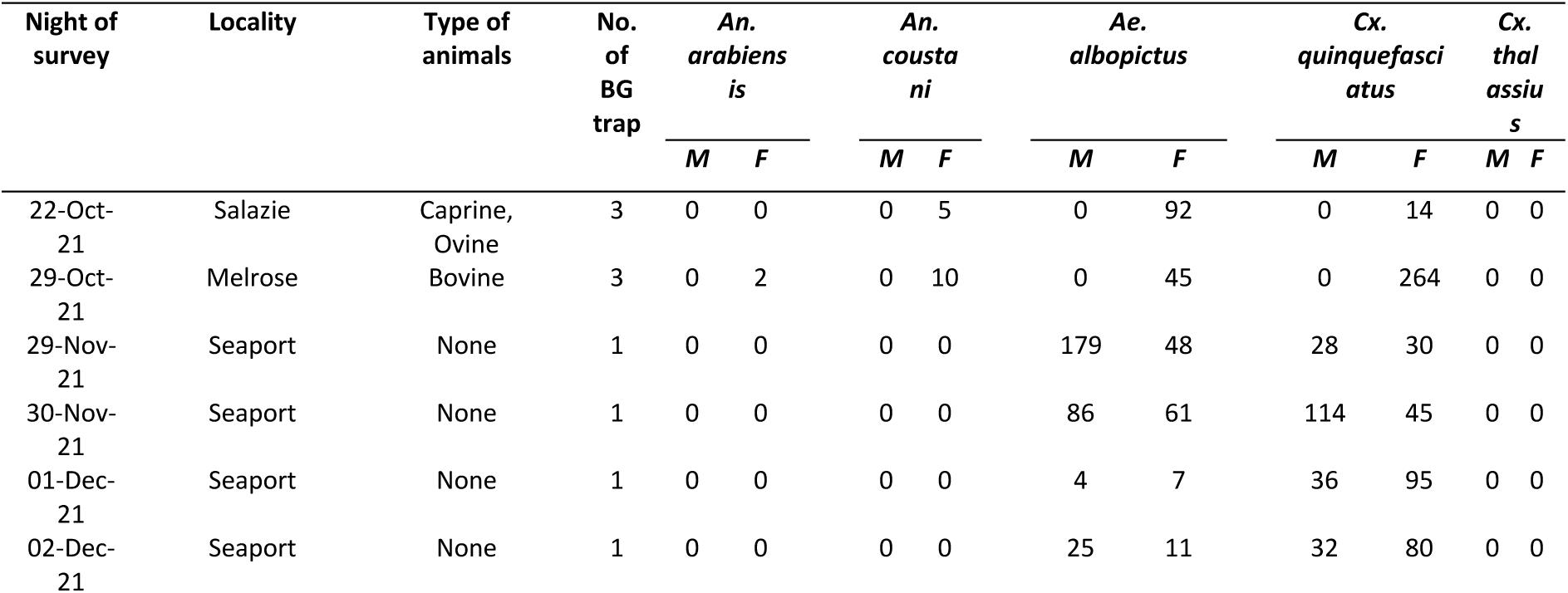
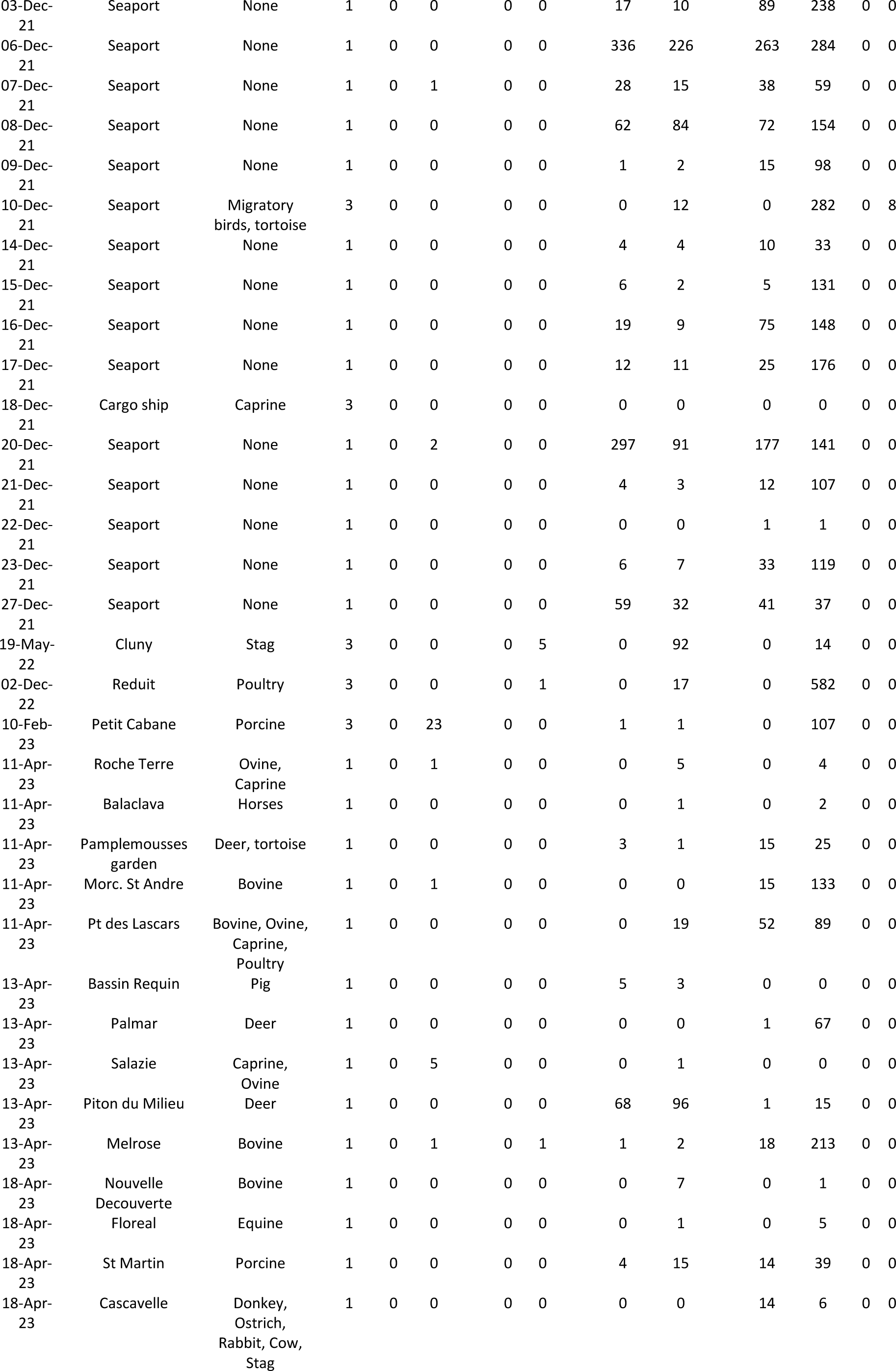
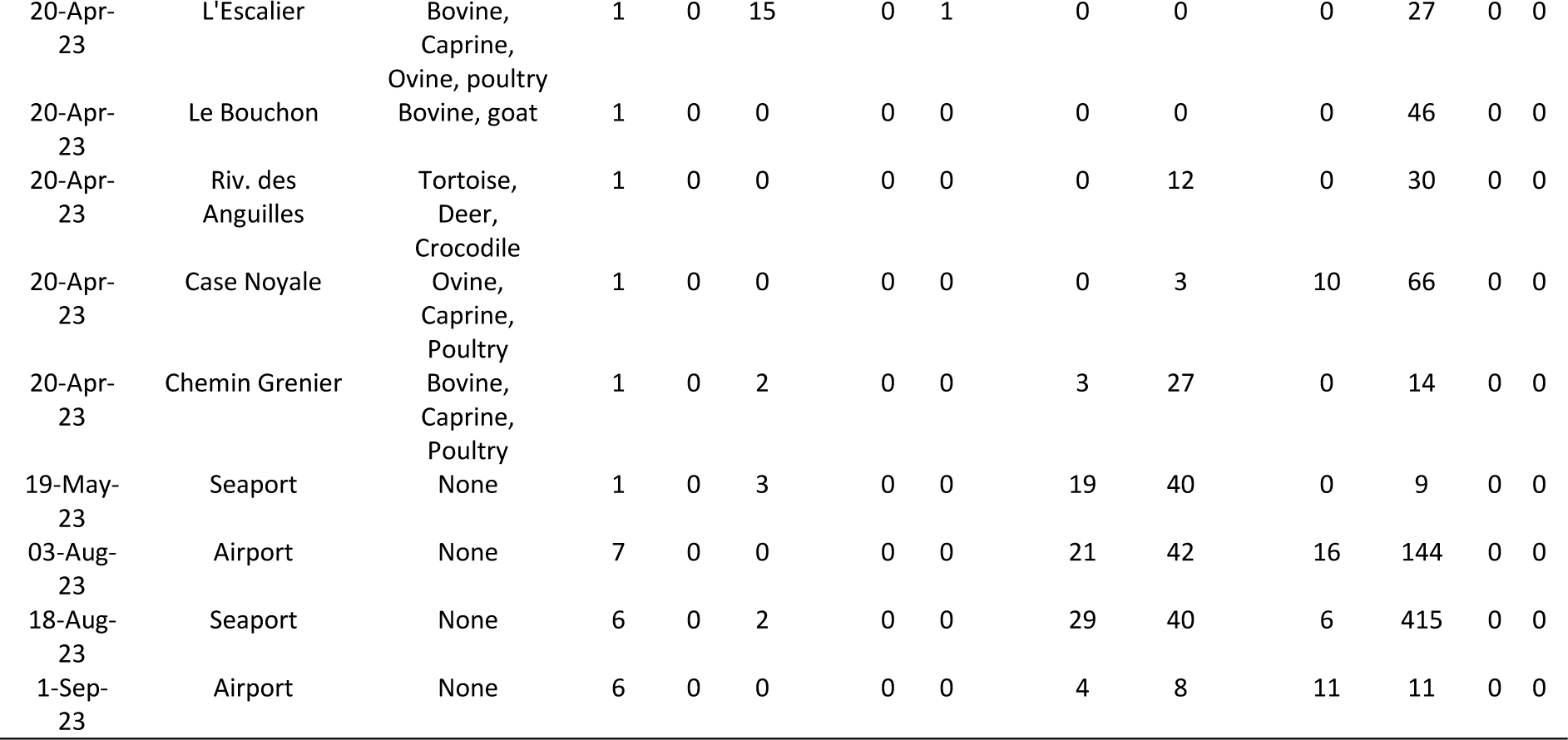
Total number of adult mosquitoes collected by BG Sentinel traps at the seaport, airport and 26 animal assembly points in Mauritius during snapshot surveys between October 2021 and September 2023.

#### 3.1.3 Surveillance of artificial mosquito breeding sites at seaport

Larvae of *Ae. albopictus, An. arabiensis, Cx. quinquefasciatus* and *Lutzia tigripes*, formerly known as *Culex (Lutzia) tigripes*, were detected in artificial breeding sites set up in the seaport area from July 2022 to April 2023 (Fig. 3). *Aedes albopictus* was highly prevalent, breeding in nearly all of the 38 artificial breeding sites. In contrast, *An. arabiensis* and *Lutzia tigripes* bred exclusively in concrete ground pools in a vegetated yard close to the disembarkation zone. *Culex quinquefasciatus* bred in two other sites besides the ground pools. No *An. stephensi* larvae were detected in these artificial breeding sites at the seaport during the entire monitoring period.

**Fig. 3:**
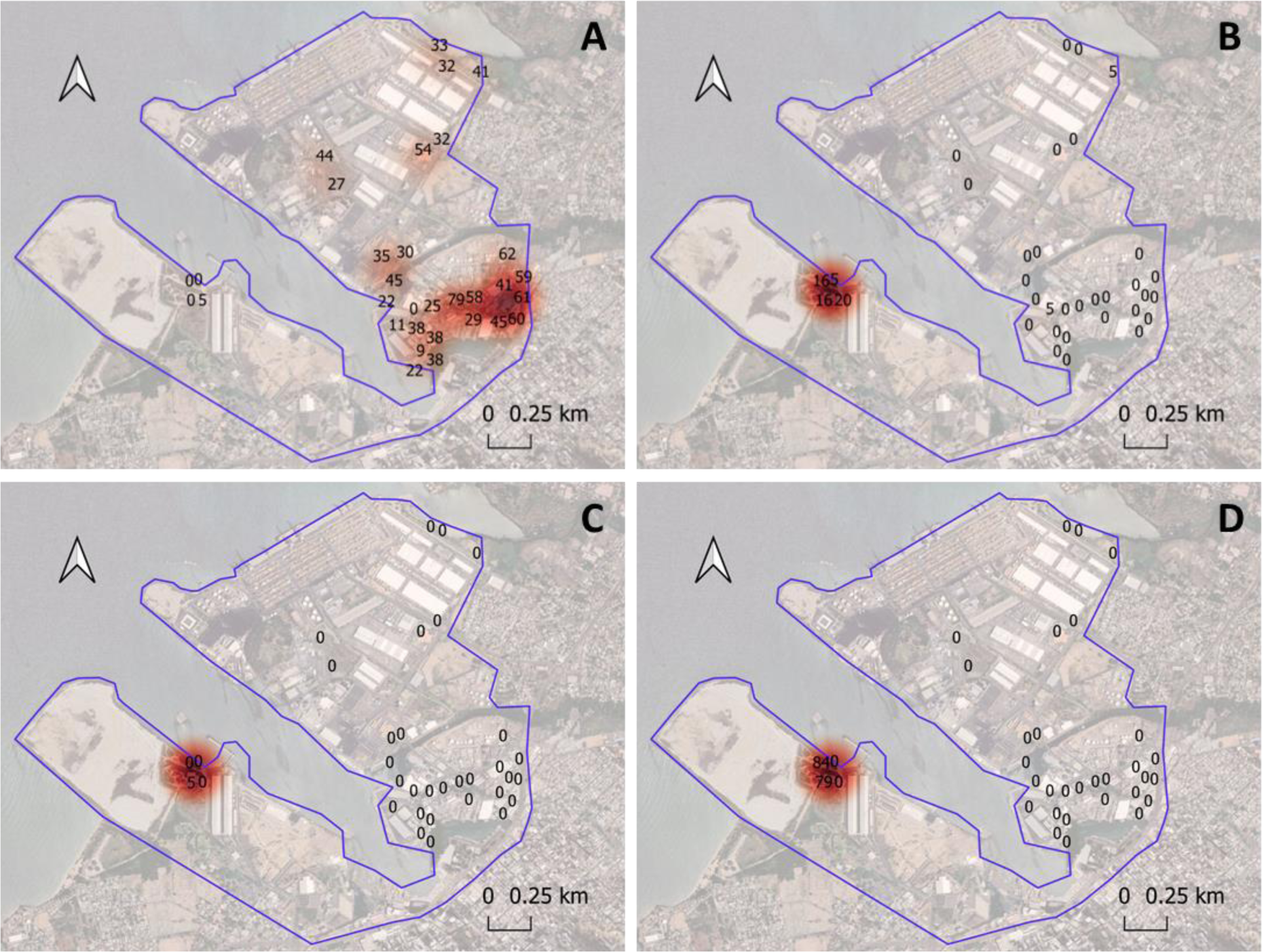
Heat maps showing frequency of artificial mosquito breeding sites positive for larvae of (**A)** *Ae. albopictus* (**B)** *Cx. quinquefasciatus*, **(C)** *An. arabiensis* and, **(D)** *Lutzia tigripes* during weekly inspections carried out from July 2022 to April 2023 in the seaport area. This figure was produced using Quantum GIS^®^ 3.16.1 software and Google Earth.

#### 3.2 Molecular identification confirmation

In order to confirm morphological identities of the *Anopheles* specimens collected, a subset of samples was sequenced and compared to database sequences for similarity. Overall, molecular analysis confirmed the absence of *An. stephensi* in Mauritius. For the 58 specimens morphologically identified as *An. gambiae sl,* five different COI haplotypes were detected. BLAST analysis of these COI haplotypes supported most morphological identifications. One haplotype carried by two specimens revealed the highest sequence similarity with *An. merus* specifically (99.83% identity score). The remaining *An. gambiae sl* haplotypes could not be distinguished to the species level using the COI sequences. Two haplotypes were detected among the *An. gambiae sl* ITS2 sequences. The ITS2 sequence BLAST revealed those unidentified *An. gambiae sl* carried the same haplotype and were *An. arabiensis* (100% identity score). The ITS2 analysis also confirmed *An. merus* identification for two specimens (100% identity score), specifically.

For the 10 specimens morphologically identified as *An. coustani,* two COI haplotypes were detected. BLAST analysis of the two haplotypes supported this designation, mostly. For COI, the highest scoring match was for *An. coustani* COI (99.67% identity, 100% sequence coverage). ITS2 sequences revealed highest scoring match for *An. coustani* (99.83% identity, 93% sequence coverage), though there was also a high level of similarity to another species (*An. ziemmani,* 99.82% identity, 87% sequence coverage).

Phylogenetic analysis of COI was used to confirm species identification and examine evolutionary relationships among *Anopheles* species found in Mauritius and globally (Fig 4 and Fig 5). *An. arabiensis* shared a clade with other members of *An. gambiae sl* in the COI tree (bootstrap = 100), while the ITS2 tree revealed sufficient support for clustering separately from other *An. gambiae* complex species (bootstrap = 93). *Anopheles merus* was distinct from other species in both the ITS2 and COI trees. In the COI tree, *An. coustani* specimens fell into a distinct clade with other members of the same species and *An. rufipes*, *An. ziemanni*, and *An. rhodesiensis* (bootstrap = 85). There was not enough sequence similarity between *An. gambiae sl* and *An. coustani* to create an ITS2 tree that included both.

**Fig 4:**
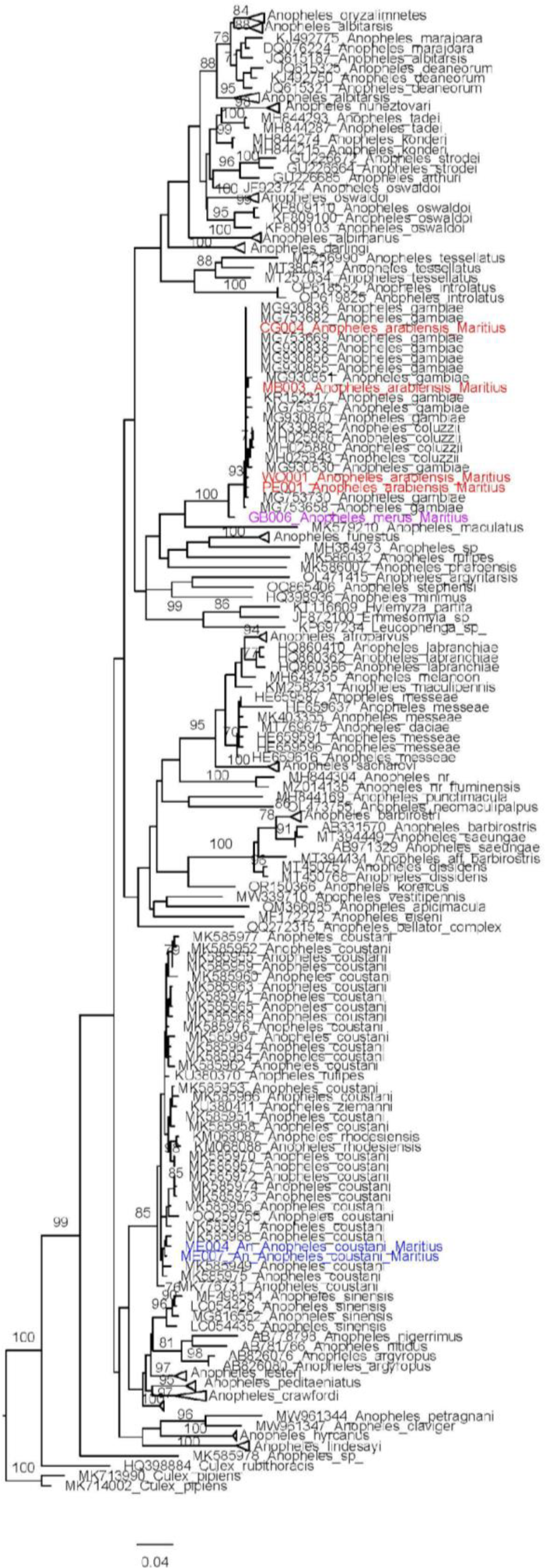
Phylogenetic analysis of COI haplotypes using maximum likelihood method. on COI haplotypes of *Anopheles* collected in Mauritius, 2021-2023, denoted in blue. Bootstrap values >70 shown at nodes.

**Fig 5:**
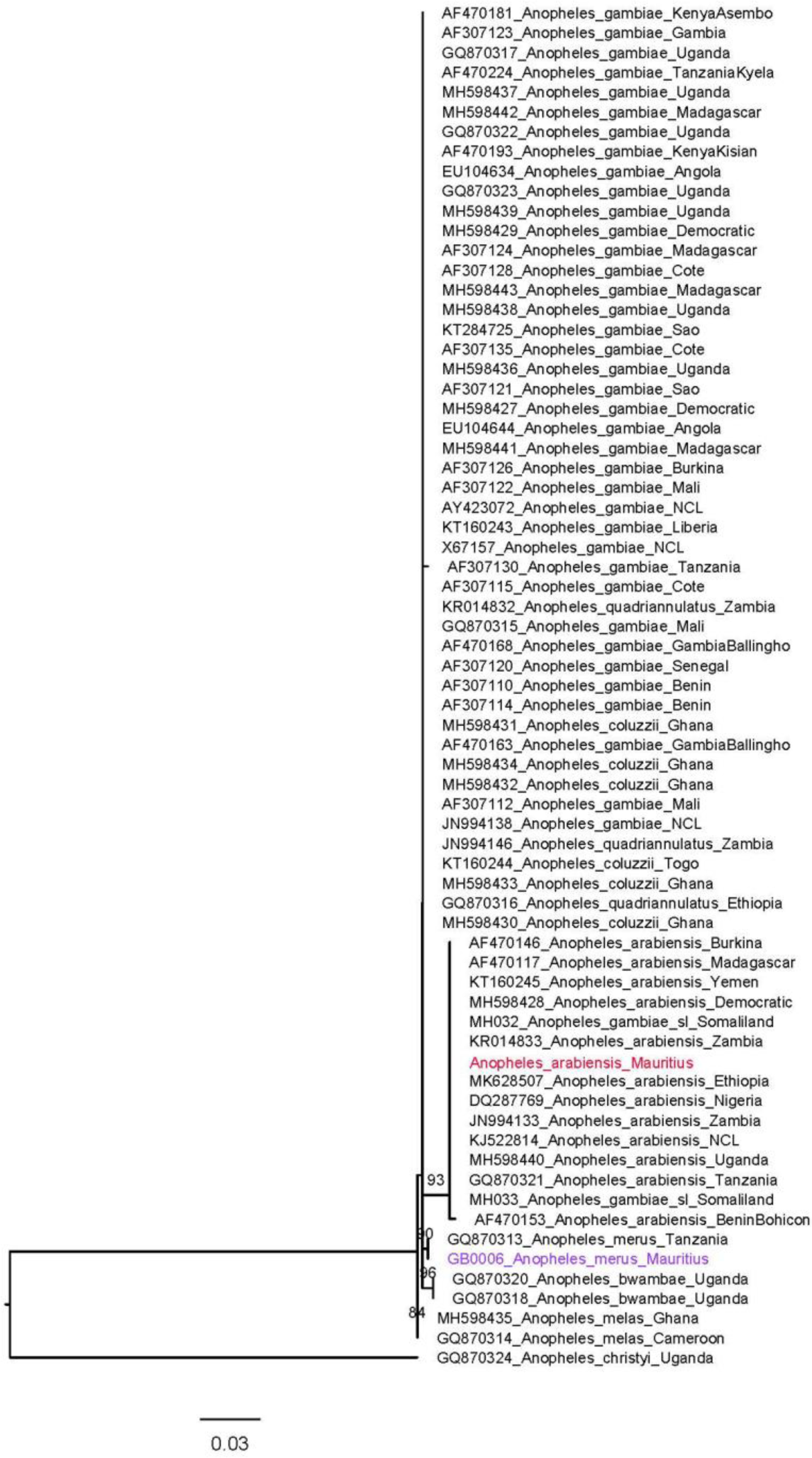
Phylogenetic analysis of ITS2 using maximum likelihood method. ITS2 haplotypes of *Anopheles* collected in Mauritius, 2021-2023, are denoted in color (purple = *An. merus*, red = *An. arabiensis*). Bootstrap values >70 shown at nodes.

## 3. Discussion

### 3.1 Main findings

Throughout the world, insect vectors and the diseases they transmit are expanding their range through international travel and trade (Liu *et al*., 2019). Environmental change (Esslf *et al*., 2019) and climate change (Huang *et al*. 2011) exacerbate the existing risks of new species (vector and/or disease agent) becoming established and existing species spreading to new areas. Island states are particularly exposed to invasive risks and the emergence of vector-borne diseases (Mavian *et al*., 2018) and integrated entomological surveillance systems aimed at detecting exotic mosquitoes and associated pathogens introduced at points of entry are routinely practised in many continental countries such as the USA (Wilke *et al*., 2022) or island countries such as New Zealand (Ammar *et al*., 2019).

By way of example, the invasive species *Ae. albopictus* has been established for several centuries in the islands of the South West Indian Ocean in connection with human population migratory episodes from Southeast Asia. In Mauritius, it was first described by de Charmoy in 1908 (de Charmoy, 1908). *Aedes albopictus* is more recently established in the Comoros, in Mayotte since 2001 (Girod, 2004), in Anjouan since 2011 (Marsden *et al*. 2013) and has just been found in Grande Comore in 2019 (Lebon, pers com). Like *Ae. aegypti*, it is a vector of numerous human arboviruses such as dengue, with the Mauritian epidemics of DENV-2 in 2009 (Issack *et al*., 2010) and more recently in 2023 (Issack, pers com), and chikungunya, with the regional epidemic in 2005-2006 (de Lamballerie *et al*., 2008). This highly competitive species is tending to out-compete *Ae. aegypti* in places where it has become established, such as Réunion (Bagny *et al*., 2009a) and Mayotte (Bagny *et al*., 2009b). In Mauritius, *Ae. aegypti* is thought to have been eliminated from the island by anti-malarial insecticide treatments carried out in the late 1940s (Halcrow, 1954).

The invasive malaria vector in Africa, *An. stephensi* was not detected in Mauritius during this 26-months period. Species collected were *Ae. albopictus*, *An. arabiensis*, *An. coustani*, *An. merus*, *Cx. quinquefasciatus*, *Cx. thalassius* and *Lutzia tigripes*, i.e. a total of 7 species, classically the most abundant of the 18 mosquito species recorded on the island (Boussès *et al*., 2018). Over and above their respective abundance in the island’s different environments (the two most ubiquitous species being *Ae. albopictus* and *Cx quinquefasciatus*), the type of collection method used greatly influenced the abundance and diversity of species collected. While *Ae. albopictus* was collected by all the three collection methods, it was the most dominant species detected during larval inspection of natural and artificial breeding sites and the second most abundant species (after *Cx. quinquefasciatus*) collected by BG Sentinel traps. *An. arabiensis*, *An. coustani, Cx. quinquefasciatus* and *Cx. thalassius* were mostly collected by BG Sentinel traps while *Lutzia tigripes* was exclusively detected in artificial concrete ground pools. This highlights the importance of using several mosquito collection methods as part of an integrated entomological surveillance, particularly when assessing the risk of introduction of invasive vector and disease transmission at points of entry (WHO, 2016).

Though *An. stephensi* was not detected in Mauritius, the inclusion of molecular analysis proved useful in this study. Using DNA sequencing for conformation, no false morphological identifications of *An. stephensi* were reported in the 2021 collection providing initial support for successful integration of *An. stephensi* morphological identification. A baseline for detection is now established should *An. stephensi* emerge. Furthermore, phylogenetic analysis of the *Anopheles* sequences provided species-level identification of *An. gambiae* specimens, confirming that two members of the *An. gambiae* complex are present in Mauritius as previously reported (Bruce-chwatt, 1974). Sequencing confirmed *An. arabiensis* for the majority of the *An. gambiae sl* collected as expected. *Anopheles merus*, a saltwater mosquito and malaria vector present in coastal regions of East Africa and Southern Africa such as Kenya, Madagascar, Mozambique, South Africa, and Tanzania, was also identified in this study (Bartilol *et al*., 2021). This vector was first identified in Mauritius in 1963 (Bruce-chwatt, 1974). While previously considered a secondary vector, studies have found *An. merus* playing a key role in malaria transmission in some areas (reviewed in Baritol et al., 2021). In addition, there is evidence that it has undergone a geographical range expansion in South Africa (Mbokazi *et al*., 2018). Thus, this vector should be monitored further.

Analysis of *An. coustani* COI and ITS2 sequences did confirm this general identification while highlighting potential challenges with distinguishing it from genetically similar species. Phylogenetic analysis of the COI sequences confirmed the distinction with *An. rhodesiensis* but there is not enough support to separate the *An. coustani* sequences from similar *An. ziemanni* and *An. rufipes* sequences included from the NCBI database. Whole genome sequencing can provide further confirmation of the identification *An. coustani* and the absence in Mauritius of other closely related species.

At both the seaport and airport, small containers, saucers of flower pots, drums, pails and buckets were the most abundant sites with water and were conducive to the breeding of *Ae. albopictus,* while the most prolific breeding site for the species were used tyres. The abundance of those types of breeding sites as well as the high prevalence of *Ae. albopictus* in nearly all of the surveyed localities, indicate the high probability of a successful colonization of *An. stephensi* if it is introduced on the island, since like *Ae. albopictus, An. stephensi* is often a container-breeder which oviposits in all kind of water and thrives in urbanized settings (Walker and Lynch, 2007; Hume *et al*., 2007; Tadesse *et al*., 2019; Fazeli-Dinan *et al*., 2022). Vector surveillance at the point of entry must target dispersal routes on a global and continental scale, but also on a local scale (Swan *et al*. 2022), taking into account the bio-ecology of the invasive species. Modelling using mainly abiotic (climate, photoperiod) and landscape (urbanisation) variables can help identify the most suitable areas for invasive species establishment. The spatial distribution of each mosquito species in the seaport area was established through the setting up and weekly surveillance of artificial breeding sites. Several hotspots for *Ae. albopictus* and therefore potentially for *An. stephensi*, have thus been identified and could be used to implement vector control response against the invasion and spread of this species in the event of early detection at this maritime entry point, the most at-risk for the island.

### 3.2 A One Health multisectoral approach

A One Health multisectoral approach was chosen to develop surveillance and rapid response strategies against a potential invasion by *An. stephensi*. The invasion of Africa by *An. stephensi* and the re-emergence of malaria in this region, particularly in Djibouti, call for the implementation of a multisectoral approach and integrated surveillance and control that should target all vectors (Al-Eryani *et al*., 2023). At a local scale, on the highly exposed island of Mauritius, the VBCD built partnerships with multiple stakeholders to strengthen vector surveillance, taking into account this new invasive risk, and to potentially intervene in case of an invasion. This is a crucial step that saves precious time during the implementation of a rapid response strategy. The Mauritian experience shows that continuous intervention, strong leadership and substantial and predictable funding are essential to prevent the re-emergence of malaria (Tatarsky *et al*., 2011). Sustained vigilance is essential given the favourable conditions in Mauritius e.g., tropical climate, *An. arabiensis* populations, commerce connectivity. The VBCD’s operational responsiveness to the risk of expansion of the new vector currently invading Africa is to be commended. In addition to this responsiveness, the multisectoral approach was initially adopted: collaborations were first established with the national Livestock and Veterinary Division (LVD) and the Mauritius Port Authority to establish an enhanced surveillance system targeting *An. stephensi* in the seaport and on cargo ships transporting livestock. Collaboration with the LVD further expanded, creating new entomological surveillance efforts for other mosquito species (*Anopheles* and *Culex* spp.) and biting midges (*Culicoides* spp.) to assess the risk of transmission of diseases of public health and veterinary interest in farms, quarantine centres, natural reserves, migratory bird areas and other animal assembly points. This in turn necessitated collaborations with the National Parks and Conservation Services, private farmers and land owners.

Moreover, establishing a surveillance system for *An. stephensi* at the airport required the collaboration of various stakeholders, including inhabitants living near the airport, the Civil Aviation Department, Airport of Mauritius Co. Ltd., Airport Terminal Operations Ltd., Beachcomber Ltd., Plaisance Meteorological Services, and two cargo companies involved with animal transport – Plaisance Air Transport Services and Gound2Air Ltd. Mosquito surveys were also conducted islandwide by the VBCD with the collaboration of the community to collect data on mosquito larval habitats. *Anopheles* (151) and *Aedes* (339) mosquitoes collected during those surveys were sent to Baylor University, Texas for PCR characterization which led to the identification of *Anopheles* spp. diversity and *Ae. albopictus* population genetics on the island. Going forward, integration of molecular surveillance as part of vector surveillance will be important for continued monitoring of vector species in Mauritius, particularly distinguishing closely related species that may be morphologically similar or unidentifiable.

Beyond mosquitoes, other invasive arthropod vectors are threatening the region. In 2002-2004, theileriosis (East Coast fever), a parasitic disease affecting ruminants and transmitted by the African tick *Rhipicephalus appendiculatus*, emerged for the first time on Grande Comore and led to the death of 10% of the island’s livestock. Following this emergence, an entomological study revealed the presence of the African vector tick *Rh. appendiculatus* in cattle herds on Grande Comoros but not yet on the other islands of the archipelago, this tick having probably been introduced to Grande Comoros by sea transport of live animals from East Africa (Yssouf *et al*., 2011). The risk of its spread to other islands in the archipelago, and even to the SWIO region, is a priority, hence the importance of setting up a genuine One Health entomological surveillance system dedicated to the vector risk to animal and human health, including the surveillance of vector-borne pathogens.

To mitigate future invasions by hematophagous arthropods and related new epidemics, it is therefore urgent to strengthen cross-border, multi-actor and multi-sector surveillance measures involving experts from the health sector, environmental stakeholders and political decision-makers, while taking account of wider social and economic perceptions.

Proactive approaches are recommended, including the operational use of new surveillance tools, such as xenomonitoring based on saliva analysis (Flies *et al*., 2015) or faeces analysis (Meyer *et al*., 2019) of mosquitoes collected in integrated trapping systems, or the development of diagnostics based on e-DNA (Kristan *et al*., 2023), or artificial intelligence algorithms for automated species recognition (e.g., Minakshi *et al*., 2020ab). Genotyping using new population genomics techniques (as high-resolution genetic markers, single nucleotide polymorphisms (SNPs)) can be undertaken to identify the origin of insects and dispersal pathways by comparing the genotype of incursion samples to reference samples of known origin (Schmidt *et al*., 2021).

Another recent development to improve the capacity to detect and monitor the spread of an invasive mosquito within a country is citizen science, where members of the public actively contribute to surveillance. Citizen science has the potential to be highly scalable with multiple collectors and the proven capacity to operate as a second-line mosquito surveillance, i.e., beyond the point of entry (Sousa *et al*. 2022; Palmer *et al*. 2017; Eritja *et al*., 2019), and as a complementary tool to existing entomological surveillance (Pernat *et al*., 2020; Sousa *et al*. 2020). Recently, several global citizen science mosquito monitoring platforms have been integrated together and with artificial intelligence (Uelmen *et al* in revision; Carney *et al*. 2022; mosquitodashboard.org), and focused specifically on the problem of *An. stephensi* invasion in Africa (Carney *et al*. 2023) and Madagascar in particular (Carney *et al*. in prep). In the present Mauritius study, the school community was encouraged to use GLOBE Mosquito Habitat Mapper, an app-based tool developed by NASA that helps the community to take an active part in surveillance by documenting mosquito breeding habitats and identifying larval mosquito genera (Low, 2018; Low *et al*., 2021; 2022). Special workshops and sensitization campaigns will be organized by the Health Ministry for a greater outreach of this tool to the Mauritian citizens. The iNaturalist platform was also utilized, and a mosquito campaign (mosquitoesInAfrica.org) was promoted locally with flyers.

### 3.3 An Indian Ocean regional approach

While genetic evidence of *An. stephensi* has not been detected in the island states and territories of the South West Indian ocean region, these countries exhibit substantial differences in malaria burden (WHO, 2022) and have huge disparities in their ability to detect and respond to a potential invasion by a new vector. For instance, malaria is still endemic in Madagascar and the Comoros; it has been eliminated in Mauritius and La Reunion, and has never been present in Seychelles due to the absence of *Anopheles* (Robert *et al*., 2011).

However, because of close economic ties among countries of the region and with the Asian and African continents, the risk of invasion by *An. stephensi* and the potential flaring up of malaria cases in one member state, may have a compounding impact on Mauritius and other neighbouring countries, potentially reversing malaria control and elimination efforts of a country. The South West Indian Ocean has historically been a hub for trade, transport and migration. As a result, countries in the region, particularly island states, share common public health threats. Among these, vector-borne and in particular mosquito-borne diseases, are prime candidates to (re)emerge and likely to spread throughout the region. The regional epidemics of chikungunya in 2005-2006 (Kariuki *et al*., 2008) and dengue fever since 2018 (Sissoko *et al*., 2009a) as well as outbreaks of Rift Valley Fever affecting the Comoros in 2007-2008 (Sissoko *et al*., 2009b) and 2019 (Roger *et al*., 2014) are good examples of this. Improving preparedness and capacity to respond to these threats at regional level is therefore a major challenge.

The Indian Ocean Commission (IOC), in the framework of its SEGA-One Health network, within the thematic pole “vector and vector-borne diseases”, has therefore been alerted to the risk of invasion by *An. stephensi* and on the need to develop a regional approach against this threat, in particular by building capacity and providing guidelines on the design and implementation of effective and sustainable entomological surveillance systems in order to improve preparedness and response. This led to the organization of a regional vector coordination workshop in June 2021, bringing together veterinarians, entomologists and public health officials from Mauritius, Réunion Island, Comoros, Seychelles, Madagascar, and partners such as the French Agricultural Research Centre for International Development (CIRAD) and “Institut Pasteur de Madagascar” (IPM). The topic continued to be discussed during the quarterly teleconference of information and experience sharing, and during the regional technical meeting of the SEGA-One Health network During those workshops and exchanges, each member state submitted a statement of vector surveillance and control capacity and gaps were identified. A regional action plan has been established, with three pillars: (i) operationalize/strengthen the vector surveillance in each member state, with a priority at the points of entry, (ii) reinforce the vector control capacity at regional level and (iii) follow up the resistance of vector to insecticide to guide vector control. As usual in the SEGA-One Health network, this regional plan involves the two sectors, veterinary services and public health authorities, to capitalize on the complementarity of the two sectors which are lab capacity and their presence in the field, including at points of entry. Practical sessions in the field and in the laboratory enabled the VBCD experience to be shared with the technical surveillance players in the region’s countries/territories. It’s important to understand that there is no one-size-fits-all entomological surveillance system. Harmonisation involves defining evidence-based standards in order to promote best practices and identify the most appropriate surveillance activities while optimising financial and human resources. Since June 2021, two workshops have been organised by the IOC with technical support from IPM and CIRAD to train two medical entomologists and two veterinary entomologists from each member state in the morphological identification of *An. stephensi*, as well as other mosquitoes, ticks, fleas and *Culicoides* species with a potential for invasion in the region. Pinned specimens of *An. stephensi* used during those training workshops were provided by the US Centers for Disease Control and Prevention (CDC). At the request of each member state, field equipment and consumables (traps, thermal foggers, larviciding apparatus, etc.) were procured and expert missions were organised by the IOC to initiate or strengthen the surveillance and potential control of *An. stephensi* on the islands. A regional workshop to facilitate the development and implementation of a regional rapid response strategy, should *An. stephensi* be detected in one of the member states, is scheduled. Another workshop focusing on a table simulation exercise in case of introduction is also in the pipeline. Although control measures are not part of the surveillance process, they are both closely linked. There is therefore an urgent need to develop response capabilities, particularly in terms of vector control, in parallel with the optimisation of surveillance systems, in line with the Integrated Vector Management (IVM) framework promoted by the WHO. Collaboration between CIRAD further developed, leading to technical and logistic assistance in the monitoring and characterization of insecticide resistance within mosquito populations of the region. Most vector control programmes rely heavily on the use of chemical insecticides, so monitoring the susceptibility of vectors to commonly used active substances should be a key component of entomological surveillance systems and an integral part of these systems. Pending the generation and analysis of data, support of the IOC has already been secured to investigate alternative products/methods to better control resistant mosquito populations (and potentially *An. stephensi*) in countries of the region. Collaboration with CIRAD and Mauritius has further developed, leading to technology transfer and capacity building of VBCD and LVD staff by two *Culicoides* specialists. Currently, a study to assess the diversity, abundance, spatial distribution and seasonal dynamics of *Culicoides* in Mauritius is currently being carried out.

## 4. Conclusion

Risk of *An. stephensi* invasion in Mauritius has triggered the Ministry of Health to initiate collaboration with partners at the national, regional and international level in order to strengthen its mosquito surveillance system. This has led to further collaboration in other fields, including initiating/strengthening surveillance of other vectors of public health and veterinary importance and characterization of insecticide resistance in the local mosquito populations. The existence of the SEGA One-Health network and strong partnerships in the area (CIRAD, IPM) represent a major asset to consider and manage the risk at national and regional levels, taking into account the risk shared by the Indian ocean islands.

In this study, mosquito larval surveys were conducted at points of entry and island-wide and adults were collected by BG Sentinel traps in the seaport, airport and at animal assembly points. Artificial breeding sites were also set up and monitored weekly in the seaport. Our experimental design was appropriate to detect important mosquito vector species that should be considered by mosquito control strategies. *An. stephensi* was not detected while *Ae. albopictus* and *Cx. quinquefasciatus* were the most prevalent species. Other species collected were *An. arabiensis*, *An. coustani*, *Cx. thalassius* and *Lutzia tigripes.* In the seaport, the diversity of mosquito species collected and their abundance were influenced by the type of collection method used - highlighting the need to use different mosquito surveillance tools to increase the likelihood of spotting an invasive species at the ports of entry.

The favorable climate, the presence of conducive larval breeding sites at the ports of entry and the high prevalence of *Ae. albopictus* (a proxy for *An. stephensi*) island-wide, indicate the high probability of a successful colonization of *An. stephensi* if it is introduced on the island – hence the need for continued surveillance efforts to a potential early detection at points of entry, and although complex due to trade and commerce regulations, potential interventions targeting disinsecticization on ships coming from countries with *An. stephensi* could be considered.

In Mauritius, as in other countries, the prevention and management of vector-borne diseases must be addressed in the prism of a "One Health" strategy that includes entomological surveillance as an integral part of the policy. Surveillance of multiple vectors at points of entry, involving all stakeholders, is a priority in veterinary and medical health, not only to inform control actions for more effective control strategies aimed at reducing populations of vector species in these areas and reducing the risk of introducing pathogens (e.g., arboviruses), but also to enable the early detection and elimination of invasive vector species inadvertently transported by recreational craft and cargo ships. Such surveillance efforts can be complemented by leveraging existing citizen science infrastructure for the detection and monitoring of invasive and vector mosquitoes, the promotion of which can in turn provide concomitant benefits to the public (e.g., awareness of *An. stephensi* and the need to remove standing water). Finally, an active regional network is needed to exchange and standardise methods and approaches in the form of guidelines, build skills and capacity through dedicated training and scientific exchanges, strengthen collaborations between disciplines and sectors at both local and regional levels, and seamlessly share effective knowledge and information between countries (i.e., online cloud-based systems) on vector surveillance and the detection of invasive insects in new areas.

## Disclaimer

The authors declare no competing interests. The findings and conclusions expressed herein are those of the authors and do not necessarily represent the official position of the U.S. Centers for Disease Control and Prevention (CDC).

## Acknowledgments

This work was funded by the Ministry of Health and Wellness, Mauritius, as part of its endeavor to improve the national mosquito surveillance system in Mauritius. Article processing charge was funded US CDC. Molecular analysis was funded through Baylor University and the US Centers for Disease Control and Prevention contract with Dr. Carter. The authors would like to thank field officers of the Vector Biology and Control Division, Ministry of Health and Wellness, Mauritius and veterinary officers of the Livestock and Veterinary Services, Ministry of Agro Industry, Mauritius, for their participation in mosquito surveys. We also extend our sincere gratitude to Dr. Russanne Low for introducing the NASA’s GLOBE Observer Mosquito Habitat Mapper to local staff.

## Author Contributions

Conceptualization: Diana P. Iyaloo, Sarah Zohdy, Tamar E. Carter, Ambicadutt Bheecarry

Data curation: Diana P. Iyaloo, Varina Ramdonee Mosawa, Tamar E. Carter, Joseph Spear, Ryan Carney

Formal analysis: Diana P. Iyaloo, Tamar E. Carter, Joseph Spear, Ryan Carney

Methodology: Diana P. Iyaloo, Varina Ramdonee Mosawa, Khouaildi Elahee, Sarah Zohdy, Tamar E. Carter, Ryan Carney

Investigation: Varina Ramdonee Mosawa, Khouaildi Elahee, Nabiihah Munglee, Nilesh Latchooman, Surendra Puryag, Hemant Bhoobun, Tamar E. Carter, Joseph Spear, Ryan Carney

Supervision: Diana P. Iyaloo, Hemant Bhoobun, Ambicadutt Bheecarry

Writing – original draft: Diana P. Iyaloo, Sarah Zohdy, Thierry Baldet

Writing – review & editing: Diana P. Iyaloo, Sarah Zohdy, Varina Ramdonee Mosawa, Khouaildi Elahee, Nabiihah Munglee, Nilesh Latchooman, Surendra Puryag, Ambicadutt Bheecarry, Hemant Bhoobun, Harena Rasamoelina-Andriamanivo, Said Ahmed Bedja, Tamar E. Carter, Joseph Spear, Ryan Carney, Thierry Baldet

## References

1. Ahn, J., Sinka, M., Irish, S. and Zohdy, S., 2023. Modeling marine cargo traffic to identify countries in Africa with greatest risk of invasion by Anopheles stephensi. Scientific Reports, 13(1), p.876.

2. Al-Eryani, S.M., Irish, S.R., Carter, T.E. et al. Public health impact of the spread of Anopheles stephensi in the WHO Eastern Mediterranean Region countries in Horn of Africa and Yemen: need for integrated vector surveillance and control. Malar J 22, 187 (2023). 10.1186/s12936-023-04545-y

3. Ammar SE, Mclntyre M, Swan T, Kasper J, Derraik JGB, Baker MG, Hales S. Intercepted Mosquitoes at New Zealand’s Ports of Entry, 2001 to 2018: Current Status and Future Concerns. Trop Med Infect Dis. 2019 Jul 5;4(3):101. doi: 10.3390/tropicalmed4030101. PMID: 31284464; PMCID: PMC6789606.

4. Bagny L, Delatte H, Quilici S, Fontenille D. Progressive decrease in Aedes aegypti distribution in Reunion Island since the 1900s. J Med Entomol. 2009a Nov;46(6):1541–5.

5. Bagny L, Delatte H, Elissa N, Quilici S, Fontenille D. Aedes (Diptera: Culicidae) vectors of arboviruses in Mayotte (Indian Ocean): distribution area and larval habitats. J Med Entomol. 2009b Mar;46(2):198–207.

6. Balkew, M., Mumba, P., Dengela, D., Yohannes, G., Getachew, D., Yared, S., Chibsa, S., Murphy, M., George, K., Lopez, K. and Janies, D., 2020. Geographical distribution of Anopheles stephensi in eastern Ethiopia. Parasites & vectors, 13(1), pp.1–8.

7. Balkew, M., Mumba, P., Yohannes, G., Abiy, E., Getachew, D., Yared, S., Worku, A., Gebresilassie, A., Tadesse, F.G., Gadisa, E. and Esayas, E., 2021. An update on the distribution, bionomics, and insecticide susceptibility of Anopheles stephensi in Ethiopia, 2018–2020. Malaria Journal, 20(1), pp.1–13.

8. Bartilol, B., Omedo, I., Mbogo, C., Mwangangi, J. and Rono, M.K., 2021. Bionomics and ecology of Anopheles merus along the East and Southern Africa coast. Parasites & vectors, 14, pp.1–11.

9. Boussès P, Le Goff G & Robert V, 2018. Inventaire des moustiques (Diptera : Culicidae) des îles du sud-ouest de l’océan Indien, Madagascar excepté : Une revue critique, Annales de la Société entomologique de France. 10.1080/00379271.2018.1429951

10. Bruce-Chwatt, L.J. and Bruce-Chwatt, J.M., 1974. Malaria in Mauritius--as dead as the dodo. Bulletin of the New York Academy of Medicine, 50(10), p.1069.

11. Carney RM, Mapes C, Low RD, Long A, Bowser A, Durieux D, Rivera K, Dekramanjian B, Bartumeus F, Guerrero D, et al. Integrating global citizen science platforms to enable next-generation surveillance of invasive and vector mosquitoes. Insects. 2022; 13(8):675. 10.3390/insects13080675

12. Carney, RM, Long, A, Low, RD, Zohdy, S, Palmer, JRB, Elias, P, Bartumeus, F, Njoroge, L, Muniafu, M, Uelmen, JA, Rahola, N and Chellappan, S. 2023. Citizen science as an approach for responding to the threat of *Anopheles stephensi* in Africa. *Citizen Science: Theory and Practice*, X(X): X, pp. 1–15. DOI: 10.5334/cstp.616

13. Carter, T.E., Yared, S., Getachew, D., Spear, J., Choi, S.H., Samake, J.N., Mumba, P., Dengela, D., Yohannes, G., Chibsa, S. and Murphy, M., 2021. Genetic diversity of Anopheles stephensi in Ethiopia provides insight into patterns of spread. Parasites & Vectors, 14(1), p.602.

14. Carney RM, Azam F, Rafarasoa LS, Riantsoa V, Rivera K, Bhuiyan T, Low RD, Zohdy S, Andrianjafy TM, Ramahazomanana MA, Rasolofo R, Subramani PA, & Chellappan S. Artificial intelligence and community science as a solution for enhanced global surveillance of invasive malaria mosquito *Anopheles stephensi*: Madagascar case study. In preparation.

15. Chalam, B.S., 1927. The Resistance of Anopheles Eggs to Desiccation. Indian Journal of Medical Research, 14(4).

16. Coetzee, M., 2020. Key to the females of Afrotropical Anopheles mosquitoes (Diptera: Culicidae). Malaria Journal, 19(1), pp.1–20.

17. de Charmoy D. 1908. On three new species of Culex collected during the anti-malarial campaign in Mauritius in 1908. Annals of Tropical Medicine and Parasitology. 2(3):257–264.

18. de Lamballerie X, Leroy E, Charrel RN, Ttsetsarkin K, Higgs S, Gould EA. Virol J. 2008 Feb 27;5:33. doi: 10.1186/1743-422X-5-33. Chikungunya virus adapts to tiger mosquito via evolutionary convergence: a sign of things to come?

19. Edwards FW (1941) Mosquitoes of the Ethiopian region. III. Culicinae adult and pupae. The Oxford University Press

20. Eritja, R., Ruiz-Arrondo, I., Delacour-Estrella, S., Schaffner, F., Álvarez-Chachero, J., Bengoa, M., Puig, M.Á., Melero-Alcíbar, R., Oltra, A. and Bartumeus, F., 2019. First detection of *Aedes japonicus* in Spain: an unexpected finding triggered by citizen science. Parasites & vectors, 12, pp.1–9.

21. Essl F, Dullinger S, Genovesi P, Hulme PE, Jeschke JM, Katsanevakis S, et al. A conceptual framework for range-expanding species that track human-induced environmental change. Bio Sci. 2019; 69:908–19.

22. Faulde, M.K., Rueda, L.M. and Khaireh, B.A., 2014. First record of the Asian malaria vector Anopheles stephensi and its possible role in the resurgence of malaria in Djibouti, Horn of Africa. Acta tropica, 139, pp.39–43.

23. Fazeli-Dinan M, Azarnoosh M, Özgökçe MS, Chi H, Hosseini-Vasoukolaei N, Haghi FM, et al. Global water quality changes posing threat of increasing infectious diseases, a case study on malaria vector Anopheles stephensi coping with the water pollutants using age-stage, two-sex life table method. Malar J. 2022; 21:178.

24. Flies, E.J., Toi, C., Weinstein, P., Doggett, S.L. and Williams, C.R., 2015. Converting mosquito surveillance to arbovirus surveillance with honey-baited nucleic acid preservation cards. Vector-borne and Zoonotic Diseases, 15(7), pp.397–403.

25. Focks DA. A Review of Entomological Sampling Methods and Indicators for Dengue Vectors. Special Program for Research and Training in Tropical Diseasess (TDR), UNICEF, UNDP, World Bank, World Health Organization; 2003.

26. Girod, R. 2004. First record of Aedes albopictus in Mayotte Island, Comoros Archipelago. Parasite 11:74.

27. Halcrow JG. 1954. Catalogue of the mosquitoes of Mauritius and Rodrigues. The Mauritius Institute Bulletin. 3: 234–248.

28. Hamon J (1953) Études biologique et systématique des Culicidae de l’île de La Réunion. Mem Inst Sci Madagascar. Série E IV:521–41

29. Huang D, Haack RA, Zhang R. Does global warming increase establishment rates of invasive alien species? A centurial time series analysis. PLoS ONE. 2011; 6:e24733.

30. Hume JC, Tunnicliff M, Ranford-Cartwright L, Day K. Susceptibility of Anopheles gambiae and Anopheles stephensi to tropical isolates of Plasmodium falciparum. Malar J. 2007; 6:139.

31. Issack MI, Pursem VN, Barkham TMS, Ng L-C, Inoue M, Manraj SS. Reemergence of Dengue in Mauritius. Emerg Infect Dis. 2010; 16:716–718. 10.3201/eid1604.09158

32. Kariuki Njenga M, Nderitu L, Ledermann JP, Ndirangu A, Logue CH, Kelly CHL, Sang R, Sergon K, Breiman R, Powers AM. Tracking epidemic Chikungunya virus into the Indian Ocean from East Africa. J Gen Virol. 2008 Nov;89(Pt 11):2754–2760. doi: 10.1099/vir.0.2008/005413-0. PMID: 18931072; PMCID: PMC3347796.

33. Kristan, M., Acford-Palmer, H., Campos, M.O., Collins, E.L., Phelan, J., Portwood, N.M., Pelloquin, B., Clarke, S., Lines, J., Clark, T.G. and Walker, T., 2023. Towards environmental detection, quantification, and molecular characterization of Anopheles stephensi and Aedes aegypti from experimental larval breeding sites. Scientific Reports, 13(1), p.2729.

34. Liu X, Blackburn TM, Song T, Li X, Huang C, Li Y. Risks of biological invasion on the belt and road. Curr Biol. 2019; 29:499–505

35. Low, R., 2018, December. Citizen Scientists as Community Agents of Change: GLOBE Observer Mosquito Habitat Mapper. In AGU Fall Meeting Abstracts (Vol. 2018, pp. IN22B-03).

36. Low, R., Boger, R., Nelson, P. and Kimura, M., 2021. GLOBE Mosquito Habitat Mapper citizen science data 2017–2020. GeoHealth, 5(10), p.e2021GH000436.

37. Low, R.D., Schwerin, T.G., Boger, R.A., Soeffing, C., Nelson, P.V., Bartlett, D., Ingle, P., Kimura, M. and Clark, A., 2022. Building international capacity for citizen scientist engagement in mosquito surveillance and mitigation: The GLOBE Program’s GLOBE Observer Mosquito Habitat Mapper. Insects, 13(7), p.624.

38. Macgregor, M.E. Mosquito surveys: a handbook for anti-malarial and anti-mosquito field workers. (ed., Baillière, Tindall and Cox), 118–135 (The Wellcome Bureau of Scientific Research, 1927).

39. Marsden CD, Cornel A, Lee Y, Sanford MR, Norris LC, Goodell PB, Nieman CC, Han S, Rodrigues A, Denis J, Ouledi A, Lanzaro GC. 2013. An analysis of two island groups as potential sites for trials of transgenic mosquitoes for malaria control. Evol Appl. 6(4): 706–20. doi: 10.1111/eva.12056.

40. Mavian C, Dulcey M, Munoz O, Salemi M, Vittor AY, Capua I. 2018. Islands as Hotspots for Emerging Mosquito-Borne Viruses: A One-Health Perspective. Viruses. 25;11(1). pii: E11. doi: 10.3390/v11010011.

41. Mbokazi, F., Coetzee, M., Brooke, B., Govere, J., Reid, A., Owiti, P., Kosgei, R., Zhou, S., Magagula, R., Kok, G. and Namboze, J., 2018. Changing distribution and abundance of the malaria vector in Mpumalanga Province, South Africa. Public health action, 8(1), pp.S39–S43.

42. Meyer, D.B., Ramirez, A.L., van den Hurk, A.F., Kurucz, N. and Ritchie, S.A., 2019. Development and field evaluation of a system to collect mosquito excreta for the detection of arboviruses. Journal of medical entomology, 56(4), pp.1116–1121.

43. Minakshi, M., Bharti, P., Bhuiyan, T. et al. A framework based on deep neural networks to extract anatomy of mosquitoes from images. Sci Rep 10, 13059 (2020a). 10.1038/s41598-020-69964-2

44. Minakshi M, Bharti P, McClinton III WB, Mirzakhalov J, Carney RM, & Chellappan S. 2020b. Automating the surveillance of mosquito vectors from trapped specimens using computer vision techniques. Proceedings of ACM COMPASS. p.105–115. 10.1145/3378393.3402260

45. MMS, Mauritius Meteorological Services. Climate of Mauritius [online]. Mauritius: Mauritius Meteorological Services; 2012, http://metservice.intnet.mu/climate-services/climate-of-mauritius.php.

46. MOH, Ministry of Health and Wellness, Health Statistics Unit [online]. Mauritius health statistics record 2022; 2022, https://health.govmu.org/health/wp-content/uploads/2023/08/Appendix-Health-Statistics-Report-2022-final.pdf

47. Palmer JRB, Oltra A, Collantes F, Delgado JA, Lucientes J, Delacour S, Bengoa M, Eritja R, Bartumeus F. Citizen science provides a reliable and scalable tool to track disease-carrying mosquitoes. Nat Commun. 2017;8:1–13.

48. Pernat N, Kampen H, Jeschke JM, Werner D. Citizen science versus professional data collection: comparison of approaches to mosquito monitoring in Germany. J Appl Ecol. 2020;00:1–10. 10.1111/1365-2664.13767.

49. Robert V, Rocamora G, Julienne S, Goodman SM. Malar J. 2011 Feb 8;10:31. doi: 10.1186/1475-2875-10-31. Why are anopheline mosquitoes not present in the Seychelles?

50. Roger M, Beral M, Licciardi S, Soule M, Faharoudine A, Foray C, et al. 2014. Evidence for Circulation of the Rift Valley Fever Virus among Livestock in the Union of Comoros. PLoS Negl Trop Dis. 8.

51. Rowland, M., Durrani, N., Kenward, M., Mohammed, N., Urahman, H. and Hewitt, S., 2001. Control of malaria in Pakistan by applying deltamethrin insecticide to cattle: a community-randomised tri*al*. The Lancet, 357(9271), pp.1837–1841.

52. Samarasekera, U., 2022. A missed opportunity? Anopheles stephensi in Africa. The Lancet, 400(10367), pp.1914–1915.

53. Schmidt TL, Endersby-Harshman NM, Hoffmann AA. Improving mosquito control strategies with population genomics. Trends Parasitol. 2021;37:907–21. 10.1016/j.pt.2021.05.002.

54. Sinka, M.E., Bangs, M.J., Manguin, S., Chareonviriyaphap, T., Patil, A.P., Temperley, W.H., Gething, P.W., Elyazar, I.R., Kabaria, C.W., Harbach, R.E. and Hay, S.I., 2011. The dominant Anopheles vectors of human malaria in the Asia-Pacific region: occurrence data, distribution maps and bionomic précis. Parasites & vectors, 4(1), pp.1–46.

55. Sinka, M.E., Pironon, S., Massey, N.C., Longbottom, J., Hemingway, J., Moyes, C.L. and Willis, K.J., 2020. A new malaria vector in Africa: predicting the expansion range of Anopheles stephensi and identifying the urban populations at risk. Proceedings of the National Academy of Sciences, 117(40), pp.24900–24908.

56. Sissoko D, Giry C, Gabrie P, Tarantola A, Pettinelli F, Collet L, D’Ortenzio E, Renault P, Pierre V, 2009a. Rift Valley fever, Mayotte, 2007-2008. Emerg Infect Dis. 15(4):568–70. doi: 10.3201/eid1504.081045.

57. Sissoko D, Giry C, Gabrie P, Tarantola A, Pettinelli F, Collet L, D’Ortenzio E, Renault P, Pierre V, 2009b. Rift Valley fever, Mayotte, 2007-2008. Emerg Infect Dis. 15(4):568–70. doi: 10.3201/eid1504.081045.

58. Sousa, L.B., Fricker, S.R., Doherty, S.S., Webb, C.E., Baldock, K.L. and Williams, C.R., 2020. Citizen science and smartphone e-entomology enables low-cost upscaling of mosquito surveillance. Science of the Total Environment, 704, p.135349. 10.1016/j.scitotenv.2019.135349.

59. Sousa, LB, Craig, A, Chitkara, U, Fricker, S, Webb, C, Williams, C and Baldock, K. 2022. Methodological diversity in citizen science mosquito surveillance: A scoping review. Citizen Science: Theory and Practice, 7(1): 8, pp. 1–19. DOI: 10.5334/cstp.469

60. Swan et al. Parasites & Vectors (2022) 15:303 10.1186/s13071-022-05413-5

61. Tadesse FG, Ashine T, Teka H, Esayas E, Messenger LA, Chali W, et al. Anopheles stephensi Mosquitoes as vectors of Plasmodium vivax and falciparum, Horn of Africa, 2019. Emerg Infect Dis. 2021;27:603–7.

62. Tadesse, F., Emiru, T., Getachew, D., Murphy, M., Sedda, L., Ejigu, L., Bulto, M., Byrne, I., Demisse, M., Abdo, M. and Chali, W., 2023. Anopheles stephensi is implicated in an outbreak of Plasmodium falciparum parasites that carry markers of drug and diagnostic resistance in Dire Dawa City, Ethiopia, January–July 2022.

63. Tatarsky A, Aboobakar S, Cohen JM, Gopee N, Bheecarry A, Moonasar D, et al. 2011. Preventing the Reintroduction of Malaria in Mauritius: A Programmatic and Financial Assessment. PLoS ONE 6(9):e23832.10.1371/journal.pone.0023832.

64. Uelmen JA, Clark A, Palmer J, Kohler J, Van Dyke LC, Low R, Mapes C, & Carney RM. In revision. Global Mosquito Observations Dashboard (GMOD): a user-friendly web interface fueled by citizen science to monitor invasive and vector mosquitoes. International Journal of Health Geographics. In revision

65. Walker K, Lynch M. Contributions of Anopheles larval control to malaria suppression in tropical Africa: review of achievements and potenti*al*. Med Vet Entomol. 2007;21:2– 21.

66. WHO, 1975. Manual on practical entomology in Malaria. Part II. Methods and techniques. Division of Malaria and other Parasitic Diseases.

67. WHO, Vector surveillance and control at ports, airports, and ground crossings. Geneva, Switzerland: World Health Organization, 2016. Available at: https://www.who.int/publications/i/item/vector-surveillance-and-control-at-ports-airports-and-ground-crossings.

68. WHO, World Health Organization and UNICEF, 2017. Global vector control response 2017–2030. Geneva: World Health Organization; 2017.Licence: CC BY-NC-SA 3.0 IGO. Available at: https://iris.who.int/bitstream/handle/10665/259205/9789241512978-eng.pdf?sequence=1

69. WHO, World Health Organization, 2021. *Health and climate change: country profile 2021: Mauritius* (No. WHO/HEP/ECH/CCH/21.01.06). World Health Organization.

70. WHO, World Health Organization, 2022. WHO initiative to stop the spread of Anopheles stephensi in Africa (No. WHO/UCN/GMP/2022.06). World Health Organization.

71. Wilke ABB, Vasquez C, Carvajal A, Moreno M, Petrie WD, Beier JC (2022) Mosquito surveillance in maritime entry ports in Miami-Dade County, Florida to increase preparedness and allow the early detection of invasive mosquito species. PLoS ONE 17(4): e0267224. 10.1371/journal.pone.0267224

72. Yssouf, A., Lagadec, E., Bakari, A., Foray, C., Stachurski, F., Cardinale, E., Plantard, O., Tortosa, P., 2011. Colonization of Grande Comore Island by a lineage of Rhipicephalus appendiculatus ticks. Parasit. Vectors 4, 38.

